# Stem cell-specific NF-κB is required for stem cell survival and epithelial regeneration upon intestinal damage

**DOI:** 10.1101/2025.02.04.636503

**Authors:** Aurélia Joly, Meghan Ferguson, Minjeong Shin, Edan Foley

## Abstract

Immune signals coordinate the repair of damaged epithelia by intestinal stem cells. However, it is unclear if immune pathways act autonomously within the stem cell to direct the damage response pathway. We consider this an important question, as stem cell dynamics are essential for formation and maintenance of the entire epithelium. We used *Drosophila* to determine the impact of stem cell-specific loss of NF-κB on tissue regeneration upon chemical injury. We found that loss of NF-κB enhanced cell death, impaired enterocyte renewal and increased mortality. Mechanistically, we showed that inhibition of stem cell apoptosis is essential for NF-κB-dependent maintenance of cell viability and tissue repair. Combined, our data demonstrate that stem cell-intrinsic NF-κB activity is essential for an orderly repair of damaged intestinal epithelia.

## INTRODUCTION

Intestinal Stem Cells (ISCs) generate the entire gut epithelium, a community of specialist cells that act in concert to extract luminal nutrients, present a physical barrier to noxious agents, and regulate host-microbe interactions. Immune pathways are particularly important modifiers of epithelial responses to extrinsic stressors, and errant immune signals underpin inflammatory illnesses that elevate the risk of colorectal cancer^1,2^. For example, defects in the NOD2 bacterial sensor are the greatest single genetic risk factor for Crohn’s disease^3–6^ and NOD2 signaling defects increase the risk of developing colorectal cancer^7,8^. Despite the importance of immune activity for epithelial function, we know little about ISC-intrinsic requirements for immune signals to protect from acute epithelial insults. As ISCs are critical for formation and maintenance of the epithelial barrier, we consider it important to resolve the extent to which immune activity directly impacts ISC function.

*Drosophila melanogaster* is an excellent model to characterize intestinal epithelial cell dynamics^9–11^. Like vertebrates, the fly gut is lined by a continuous epithelial layer that is maintained by basal, multipotent ISCs^12,13^. In flies and vertebrates, ISCs typically divide asymmetrically to generate a niche-resident daughter stem cell, and transient, undifferentiated cells that exit the niche and migrate apically^14–17^, where they matures as absorptive enterocytes, or specialized enteroendocrine cells^18^. In flies, the transient enterocyte precursor is classified as an enteroblast, and the stem cell-enteroblast pair constitutes the midgut progenitor compartment. Aside from similarities in cell identity and fate choices, ISC function is governed by similar signaling cassettes in flies and vertebrates. In both cases, EGF, JAK-STAT and BMP pathways control ISC proliferation, while signals from the Notch pathway promote enterocyte fate choices among progenitors^16,18–30^. Given the similarities between fly and vertebrate intestinal epithelial organization, *Drosophila* has considerable potential to uncover foundational aspects of immune regulation of epithelial stress responses.

The *Drosophila* Immune Deficiency (IMD) pathway, an antibacterial response with overt similarities to the mammalian NOD2 pathway, is a prominent regulator of fly responses to gut microbes^31–33^. Bacterial diaminopimelic-acid-containing peptidoglycan engages host pattern recognition proteins that signal through the Imd protein to activate fly IKK and NF-κB orthologs^34^. Active NF-κB modifies intestinal expression of antimicrobial peptides, stress response pathway genes, and genes that control metabolic activity^33^. Recent work uncovered cell-type specific roles for the IMD-NF-κB axis in mature epithelial cells^35,36^. In contrast, we know little about ISC-intrinsic roles for IMD in the control of stem cell responses to acute stresses, despite evidence that IMD pathway components are expressed at relatively high levels in the progenitor compartment^35,37^.

We used the fly to characterize NF-κB-dependent control of ISC responses to epithelial damage. We discovered that damage activates NF-κB in stem cells, leading to IMD-dependent promotion of ISC survival and generation of transient enteroblasts. Targeted inactivation of NF-κB in ISCs resulted in death of damaged stem cells, a failure to repair damaged tissue, and substantially impaired survival of challenged flies. Our work expands our appreciation of immune-dependent control of gut homeostasis and uncovers a stem cell-intrinsic requirement for an NF-κB damage response in the gut progenitor compartment.

## RESULTS

### NF-κB Regulates Intestinal Stem Cell Dynamics

We showed that genetic inactivation of IMD within the female midgut progenitor compartment impairs homeostatic stem cell proliferation and modifies enteroendocrine cell fate specification^35^. As IMD signals involve transient activation of the stress-associated JNK pathway, and prolonged engagement of the immune-regulatory NF-κB/Relish response^38^, we asked what effects inhibition of *Relish* (*Rel*) alone has on epithelial cell dynamics. To characterize effects of Rel deficiency on the midgut epithelium, we used the temperature controlled *esg^ts^* driver line to express a validated *Rel* RNAi^39^ construct in adult female midgut progenitors (*esg^ts^/Rel^RNAi^*). Inactivation of *Rel* did not have discernible effects on adult viability relative to wildtype (*esg^ts^/*+) controls (Supplementary Figure 1), confirming that progenitor cell IMD pathway activity is not essential for survival under laboratory conditions.

As the IMD pathway regulates epithelial cell dynamics, and gut architecture typically deteriorates with age, we also compared intestinal physiology in young and old wildtype and *esg^ts^/Rel^RNAi^*flies. We did not observe differences between *esg^ts^/+* and *esg^ts^/Rel^RNAi^* flies at early stages of adulthood. In both cases, the gut contained an orderly epithelium with evenly spaced GFP-marked progenitors interspersed among a lattice of mature enterocyte and enteroendocrine cells (Supplementary Figure 2A), as well as approximately equal incidences of ISC proliferation (Supplementary Figure 2C, day 10). Matching prior studies^40^, we found that thirty-day-old wild-type intestines displayed hallmarks of age-linked dysplasia that included irregular epithelial cell spacing (Supplementary Figure 2B), and significantly elevated numbers of proliferative stem cells (Supplementary Figure 2C, day 30). In contrast, progenitor-restricted loss of *Rel* prevented age-dependent decline of epithelial organization (Supplementary Figure 2B) and resulted in significantly fewer stem cell mitoses (Supplementary Figure 2C, d30). Progenitor-specific knockdown of the IKKγ ortholog *Kenny* (*key*) also prevented age-dependent increases in mitotic stem cells (Supplementary Figure 2D), confirming a central role for the IKK/NF-κB axis in age-associated deterioration of ISC function.

As stem cells maintain the progenitors needed to generate a mature epithelium, and progenitor-specific loss of Rel affected epithelial organization in older flies, we quantified the impact of *Rel* depletion on the number and identity of midgut progenitors in thirty-day-old flies. Consistent with links between progenitor cell immune activity and ISC proliferation, we observed significantly fewer midgut progenitors in *esg^ts^/Rel^RNAi^* flies relative to *esg^ts^/+* counterparts (Supplementary Figure 2E). To test if NF-κB activity also influences cell identity within the progenitor compartment, we quantified the stem cell to enteroblast ratio in midguts of *esg^ts^/Rel^RNAi^* flies and *esg^ts^/+* controls. To do so, we employed a line that allows progenitor-specific depletion of *Rel* while marking enteroblasts with CFP and GFP reporters, and ISCs with CFP alone^41^ (*esgGAL4 UAS-CFP, Su(H):GFP/UAS-Rel^RNAi^; GAL80^ts^*, Supplementary Figure 2F). In wild-type intestines (*esgGAL4 UAS-CFP, Su(H):GFP/+; GAL80^ts^*), roughly 50% of all progenitors expressed enteroblast markers, suggesting approximately equal numbers of stem cells and enteroblasts (Supplementary Figure 2G). In contrast, only 40% of *Rel-*deficient progenitors expressed enteroblast markers (Supplementary Figure 2F, G), indicating that loss of Rel impairs enteroblast generation. Combined, our data implicate intestinal Rel activity in homeostatic control of progenitor cell proliferation and identity.

### ISC-specific NF-κB Promotes Damage-Dependent Epithelial Regeneration

As progenitors are critical for renewal of damaged epithelia, we expanded our work to determine if compromised progenitor cell immunity also affects epithelial regeneration after a period of enteric stress. Specifically, we characterized progenitor dynamics in control *esg^ts^/+* and *esg^ts^/Rel^RNAi^* flies that we fed Dextran Sodium Sulfate (DSS), a polysaccharide that damages the epithelium, promoting a rapid regenerative response driven by proliferative expansion of the ISC pool^42,43^. As expected, DSS visibly disrupted the organization of intestinal epithelial cells in *esg^ts^/+* and *esg^ts^/Rel^RNAi^*midguts (Supplementary Figure 3A). Although highly variable from animal to animal, loss of Rel did not diminish the rates of ISC proliferation (Supplementary Figure 3B), or the generation of Delta+ stem cells throughout the posterior midgut (Supplementary Figure 3C), indicating an intact proliferative response in Rel-deficient progenitors. Nonetheless, depletion of Rel from stem cells significantly impaired the ability of flies to survive a continuous DSS challenge. Within ten days of exposure to DSS, more than 85% of all *ISC^ts^/Rel^RNAi^* flies died, whereas 60% of the control *ISC^ts^/+* population remained alive (Supplementary Figure 3D), pointing to essential roles for stem cell-specific Rel activity in navigating an enteric stress.

To test requirements for Rel in epithelial regeneration after a DSS challenge, we used the *esg^ts^*Flip-Out (*esg^ts^ F/O*) lineage tracing system to track epithelial renewal for three days after treating control or *Rel*-deficient progenitors with DSS. In brief, the *esg^ts^ F/O* system marks clones of *esg*-positive progenitors and their offspring with GFP in the adult midgut^27^. As anticipated, unchallenged flies generated a limited number of small, GFP+ wildtype (*esg^ts^ F/O*/*+*) or Rel-deficient (*esg^ts^ F/O*/ *Rel^RNAi^*) clones (Figure 1A). In contrast, we noted sharp differences in the number of DSS-challenged *esg^ts^ F/O*/*+* and *esg^ts^ F/O/Rel^RNAi^* clones. Whereas wildtype progenitors routinely generated large GFP-marked clones (Figure 1A-B), *Rel*-deficient progenitors produced very few clones that were markedly reduced in size (Figure 1A-B). Thus, it appears that progenitor-specific *Rel* activity is dispensable for progenitor cell proliferation, but essential for epithelial renewal after enteric stress.

**Figure 1.**
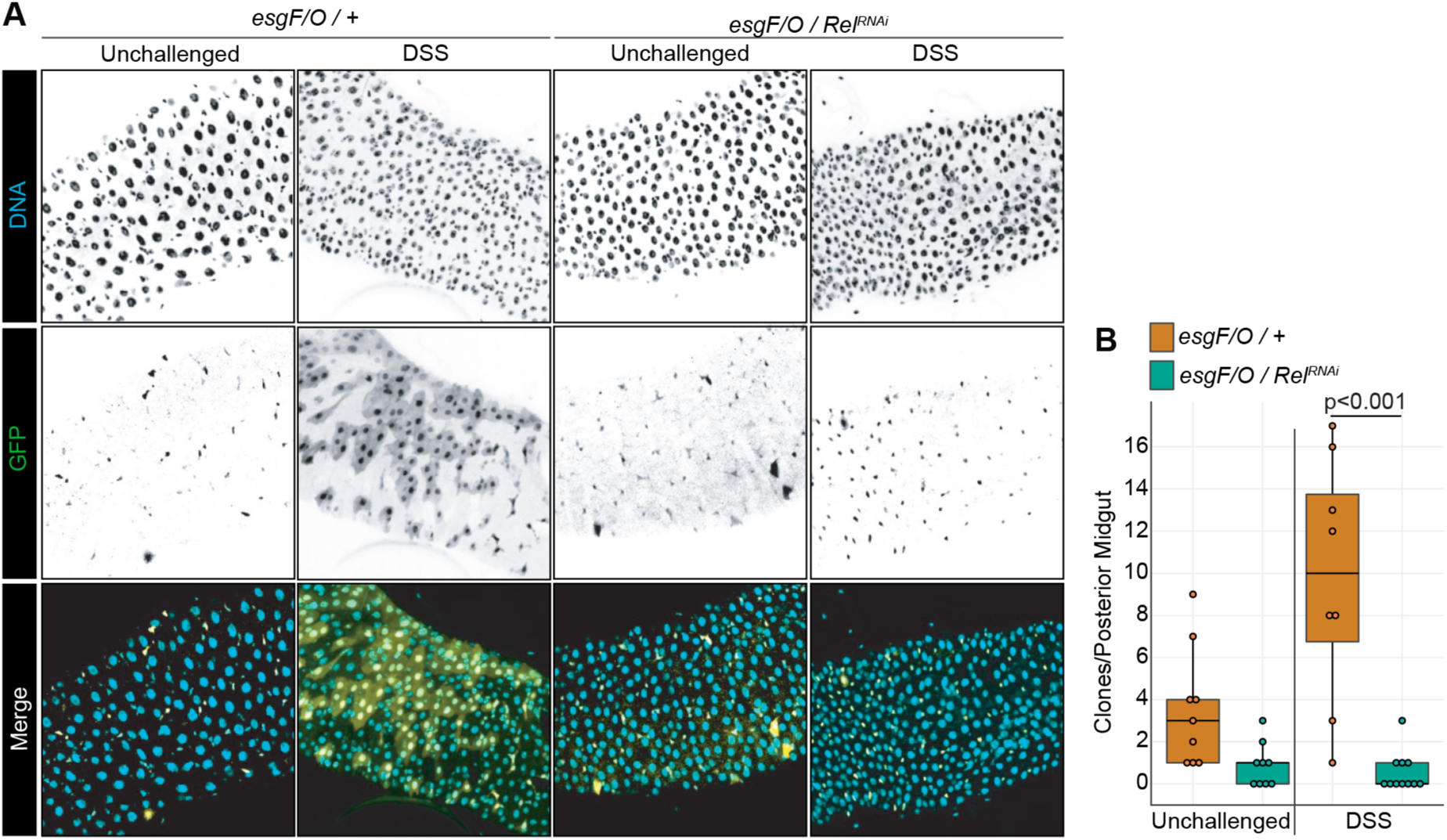
Progenitor-Specific Rel Activity is Essential for Epithelial Regeneration after a DSS challenge. **(A)** Representative images of DNA and GFP-marked clones generated by wt (*esg FO*/*+*) and Rel-deficient (*esg FO/Rel^RNAi^*) progenitors in unchallenged or DSS-treated intestines. In the merged image, DNA is visualized in cyan, and GFP-positive clones are in yellow. **(B)** Quantification of clone numbers in the posterior midguts of flies of the indicated genotypes and treatment groups. Significance was calculated using ANOVA followed by pairwise Tukey tests.

### ISC-Specific NF-κB Supports Stem Cell Survival During Enteric Stress

To better understand the involvement of stem cell-specific immunity in epithelial renewal, we performed single-cell gene expression analysis on midguts dissected from unchallenged, or DSS-treated, *ISC^ts^/+* and *ISC^ts^/Rel^RNAi^* flies. For each group, we generated transcription profiles of roughly 2,500 cells after exclusion of doublets and low-quality reads. For example, control *ISC^ts^/+* flies yielded 2,589 cells resolved in seven distinct transcriptional clusters (Supplementary Figure 4A) that expressed established markers^37^ of progenitors; pre-enteroendocrine cells; enteroendocrine cells; anterior, middle, and posterior enterocytes; and mature copper cells (Supplementary Figure 4B). We further resolved two clusters each from anterior, middle, and posterior enterocytes (Supplementary Figure 4C-D) with distinct transcription profiles for each subset (Supplementary Figure 4E). Likewise, we uncovered seven prospero-positive enteroendocrine clusters (Supplementary Figure 4F). Of the enteroendocrine clusters, cluster seven also expressed the *esg* progenitor marker, indicating that it corresponded to the committed enteroendocrine precursor (Supplementary Figure 4G), whereas the other six clusters had unique, non-overlapping peptide hormone expression profiles, suggesting that each cluster corresponds to a distinct enteroendocrine subtype (Supplementary Figure 4H).

In agreement with supplementary Figures 1 and 2, cellular identities within unchallenged ten-day-old *ISC^ts^/+* and *ISC^ts^/Rel^RNAi^* midguts were broadly similar (Supplementary Figure 5A). In both instances, we identified approximately equal ratios of progenitor, enteroendocrine, and enterocyte lineages, as well as specialist copper cells dedicated to pH regulation (Supplementary Figure 5B-C). Inactivation of Rel in stem cells had the anticipated inhibitory effect on progenitor-specific immune signals (Supplementary Figure 5D), confirming the efficacy of the *Rel^RNAi^* line. However, the consequences of Rel inactivation in stem cells extended beyond deregulated immune signals, matching earlier demonstrations that Rel has far-reaching effects on midgut transcriptional programs^32^. For example, inactivation of Rel attenuated expression of genes required for growth and development in stem cells, and well as diminished expression of genes associated with metabolism and detoxification in mature enterocytes (Supplementary Figure 5D). Loss of Rel had particularly profound effects on expression of genes linked with the stem cell to enteroblast transition (e.g. *E(spl)mgamma-HLH*, *E(spl)m3-HLH*, *Egfr*, *Dl*) within the progenitor compartment (Supplementary Figure 5E), overlapping with the loss of enteroblasts in Rel-deficient progenitors (Supplementary Figure 2). In sum, our transcriptional data indicate that we have accurately identified all prominent epithelial cell types in the *Drosophila* midgut, and our data allow us to identify specific impacts of Rel inactivation on functional states within the intestine.

**Figure 2.**
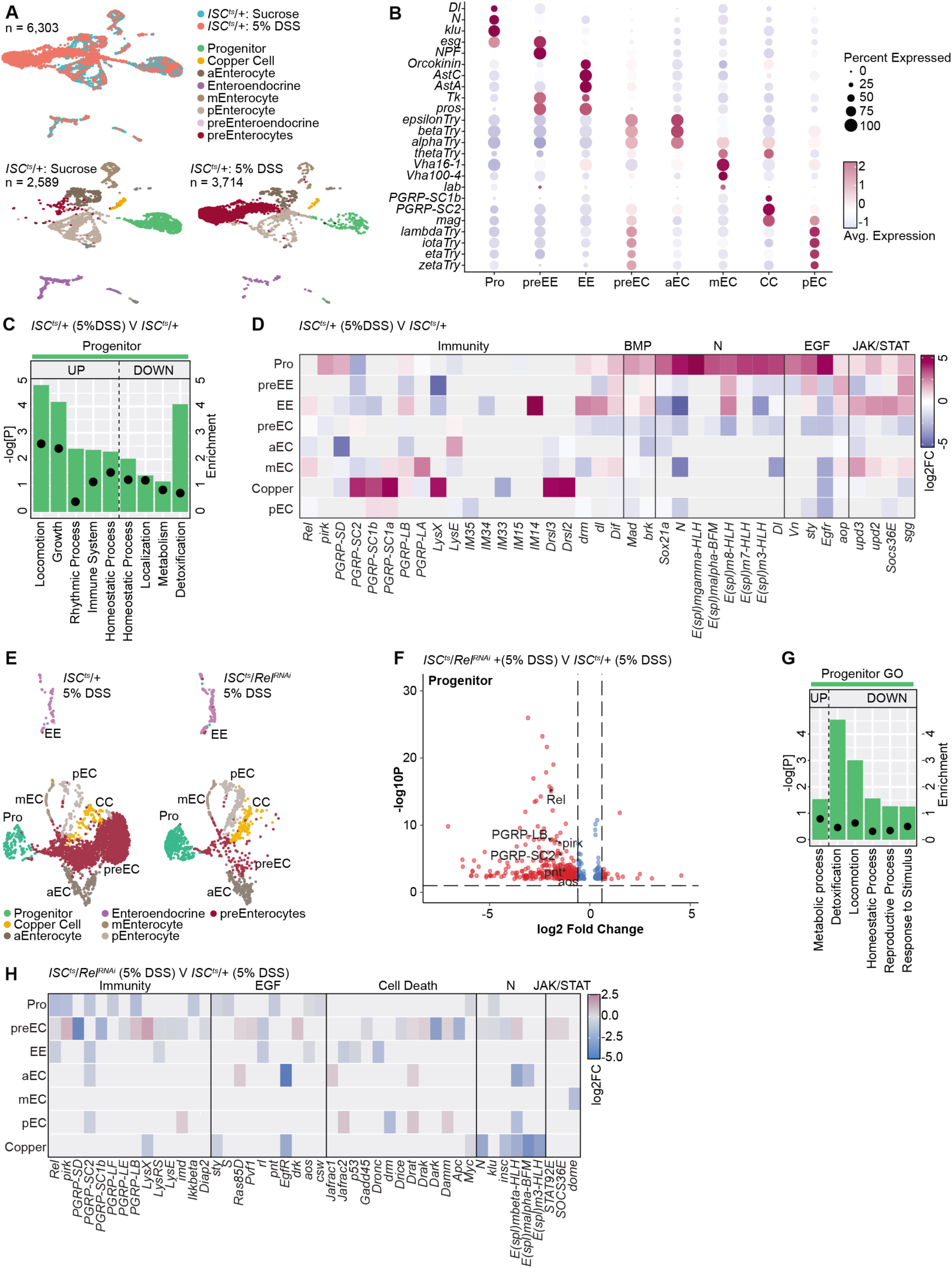
Loss of Rel in Stem Cells Modifies Expression of Apoptosis regulators in DSS-Treated Guts. **(A)** Integrated and individual UMAP projections of cell types found in midguts from ten-day-old, sucrose or DSS-treated ISCts/+ flies. **(B)** Dotplot representation of the expression levels of established markers in the indicated cell types of DSS-challenged flies. **(C)** Representation of the top five up, or downregulated gene ontology terms in DSS-exposed ISCts/+ progenitor cells relative to unchallenged ISCts/+ controls. Relative enrichment score for each term is indicated by bar length and the negative log p-values are shown with closed circles. **(D)** Heatmap illustration of representative gene expression in DSS-treated ISCts/+ midguts relative to untreated ISCts/+ controls. **(E)** Integrated and individual UMAP projections of cell types found in midguts from ten-day-old, DSS-treated ISCts/+ and ISCts/RelRNAi flies. **(F)** Volcano plot showing progenitor-specific gene expression in DSS-treated ISCts/RelRNAi midguts relative to DSS-treated ISCts/+ controls. A set of significantly modified genes are indicated. **(G)** Representation of the top five up, or downregulated gene ontology terms in DSS-exposed ISCts/RelRNAi progenitor cells relative to DSS-treated ISCts/+ controls. Relative enrichment score for each term is indicated by bar length and the negative log p-values are shown with closed circles. **(H)** Heatmap illustration of representative gene expression in DSS-treated ISCts/RelRNAi midguts relative to DSS-treated ISCts/+ controls. For each graph, pro = progenitor; preEE = pre-enteroendocrine; ee = enteroendocrine; preEC = immature enterocytes; aEC, mEC, and pEC = anterior, middle and posterior enterocytes, respectively; and CC = copper cells.

To generate an epithelium-wide map of the impacts of DSS on intestinal epithelial cell states, we next performed an integrated analysis of single cell gene expression profiles from 6,303 cells derived from unchallenged or DSS-treated *ISC^ts^/+* flies (Figure 2A). Alongside the anticipated populations of progenitor cells, and differentiated specialist cells, we found that DSS-treated wildtype guts were dominated by a cell type that is rare in unchallenged intestines (Figure 2A-B). As these cells primarily expressed low levels of anterior, middle, and posterior enterocyte markers (Figure 2B), and an earlier study showed an accumulation of immature enterocytes in DSS-treated flies^42^, we have provisionally annotated these cells as pre-enterocytes, although we caution that additional experimental data is required to determine their exact developmental trajectory. Treatment with DSS prompted extensive cell type-specific transcriptional responses within the gut that included engagement of growth and homeostatic gene ontologies within the progenitor compartment (Figure 2C). Examination of the progenitor response revealed induction of the JAK-STAT and EGF growth and repair pathways, as well as initiation of the Notch and BMP cell fate specification responses, pointing to directed epithelial regeneration within challenged intestines (Figure 2D). Alongside engagement of growth and repair pathways, we discovered that exposure to DSS also drove an immune response within progenitor cells (Figure 2C) that included transcriptional induction of Rel response genes such as *pirk*, *PGRP-SD*, and *PGRP-LB* (Figure 2D).

As DSS activates Rel in progenitors (Figure 2C-D), and Rel is essential for regeneration after enteric challenges with DSS (Figure 1), we then compared gene expression profiles of DSS-challenged *ISC^ts^/Rel^RNAi^* relative to DSS-treated *ISC^ts^/+* controls to determine what effects stem cell immune signals have on epithelial regeneration. In agreement with a requirement for Rel in epithelial renewal (Figure 1), we observed significantly fewer immature enterocytes in DSS-challenged *ISC^ts^/Rel^RNAi^*midguts relative to DSS-treated *ISC^ts^/+* controls (Figure 2E, Supplementary Figure 6). We found that blocking Rel in stem cells significantly attenuated Progenitor-specific expression of *Rel* alongside multiple IMD/Rel targets (e.g. *pirk*, *PGRP-LB*, *PGRP-SC2*, Figure 2F, H), further confirming the specificity of the *Rel^RNAi^* line. However, the impact of *Rel* inhibition extended beyond impaired immune responses and included significant inhibition of expression of genes linked to gut homeostasis and signaling (Figure 2G). In particular, we found that blocking *Rel* in stem cells attenuated expression of EGFR-Ras pathway components (Figure 2F, H), and modified expression of multiple genes linked to the regulation of cell survival and death (Figure 2H).

### Rel promotes stem cell survival during enteric stress

Our transcriptomic and functional data suggest failed epithelial regeneration and modified expression of cell survival genes in DSS-treated guts with immune-compromised stem cells. To test the hypothesis that *Rel* is necessary for stem cell survival during exposure to DSS, we used TUNEL labeling to quantify apoptotic cells in challenged and unchallenged *ISC^ts^/+* and *ISC^ts^/Rel^RNAi^* flies. Matching our earlier data, loss of *Rel* has no visible effects on gross intestinal morphology in the absence of DSS exposure (Figure 3A). However, quantification of TUNEL+ ISCs suggested a minor, statistically insignificant elevation of stem cell apoptosis in *ISC^ts^/Rel^RNAi^*midguts relative to *ISC^ts^/+* controls (Figure 3B). By contrast, loss of *Rel* greatly impacted stem cell survival in midguts exposed to DSS. On average, we observed roughly twice as many apoptotic stem cells in DSS-treated *ISC^ts^/Rel^RNAi^* midguts compared to their DSS-treated *ISC^ts^/+* counterparts (Figure 3A-B), suggesting a requirement for Rel to prevent stem cell death during exposure to DSS.

**Figure 3.**
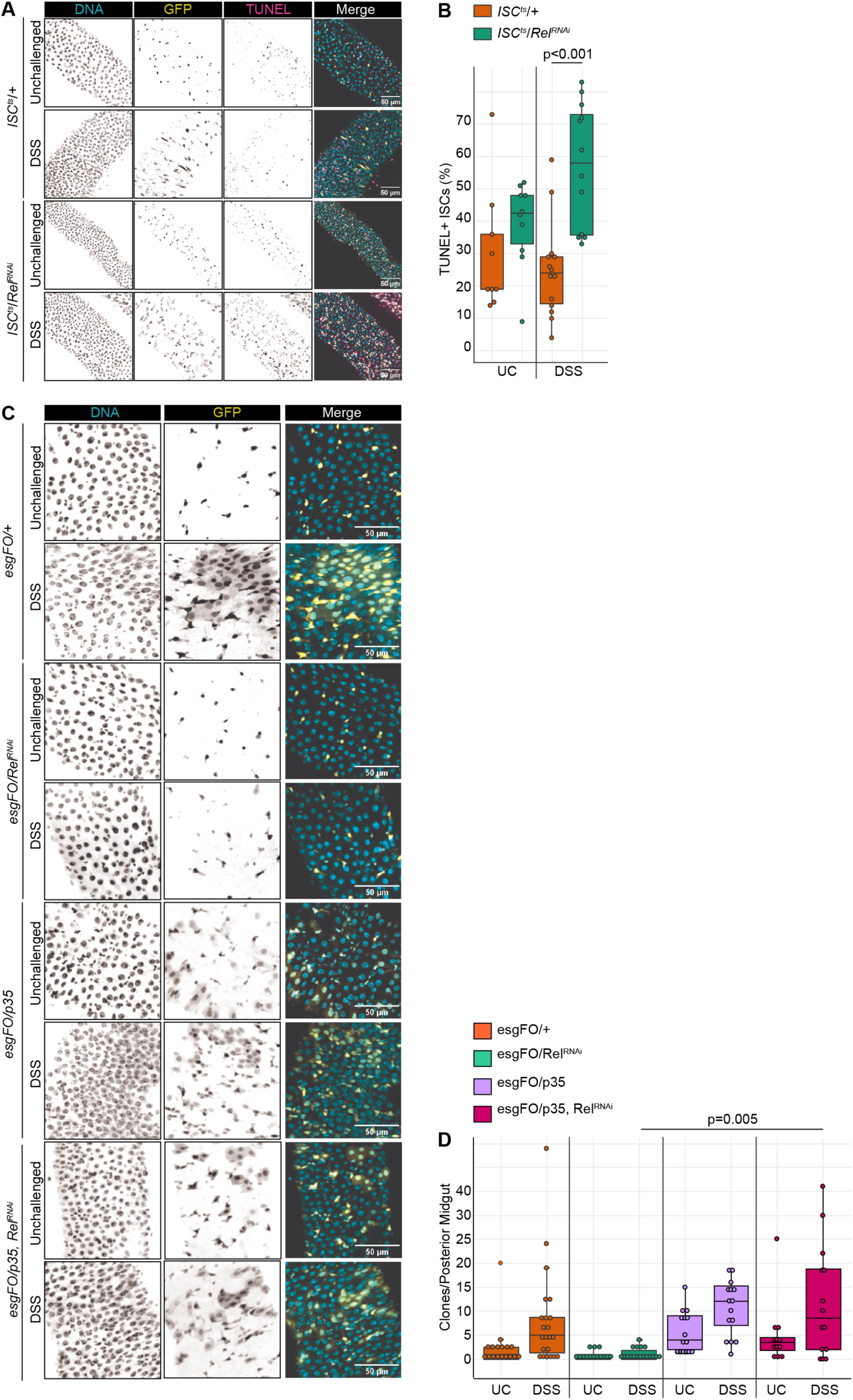
(A) Representative images of DNA, GFP-marked progenitors and TUNEL-positive cells in midguts of control or DSS-treated wildtype (*ISC^ts^*/*+*) and Rel-deficient (*ISC^ts^/Rel^RNAi^*) flies. In each merged image, DNA is cyan, GFP-positive cells are yellow, and TUNEL-positive cells are magenta. **(B)** Proportion of TUNEL positive ISCs in control, and DSS-treated *ISC^ts^/+* and *ISC^ts^/Rel^RNAi^* intestines. **(C)** Representative images of DNA and GFP-marked clones generated by unchallenged or DSS-treated intestines from flies of the indicated genotypes. In each merged image, DNA is cyan and GFP-positive clones are yellow. **(D)** Quantification of clone numbers in the posterior midguts of flies of the indicated genotypes and treatment groups. Significance for B and D were calculated using ANOVA followed by pairwise Tukey tests.

To determine if loss of immune signals prevents epithelial repair due to excess stem cell death, we used the *esg^ts^ F/O* lineage tracing system to track midgut regeneration in control flies (*esg^ts^ F/O*/*+*); flies that lack *Rel* within the progenitor compartment (*esg^ts^ F/O*/*Rel^RNAi^*); flies that express the anti-apoptotic p35 protein specifically in progenitors (*esg^ts^ F/O*/*p35*), and flies that express p35 specifically in Rel-deficient progenitors *esg^ts^ F/O*/*Rel^RNAi^, p35*). Matching our earlier work, we detected large GFP-marked clones specifically in DSS-treated wildtype flies that were absent from DSS-treated *Rel*-deficient flies (Figure 3C-D). As expected, progenitor-specific expression of p35 alone increased the number of clones produced by wildtype cells in the absence or presence of a DSS challenge (Figure 3C-D). Notably, expression of p35 was sufficient to restore epithelial regeneration to DSS-exposed Rel-deficient clones (Figure 3C-D).

In sum, our data indicate that loss of Rel impacts stress-dependent expression of apoptosis regulators and prevents the regeneration of a damaged epithelium, whereas inhibition of apoptosis within Relish-deficient cells is sufficient to restore regenerative capacity, indicating that Rel promotes stem cell survival in the *Drosophila* midgut.

## DISCUSSION

Extrinsic factors instruct the behavior of intestinal stem cells. For instance, myeloid and lymphoid cells direct ISC proliferation to match the loss of dying cells caused by microbial challenges, while germline-encoded microbial pattern recognition receptors modify ISC survival and proliferation during periods of acute, toxic stress^1,44–48^. Collectively, these observations emphasize an organ-level impact of immune activity on ISC survival and proliferation. However, the complexity of cell types within an animal gut makes it difficult to accurately resolve the effects of ISC-intrinsic immunity on gut function. We consider this an important question, as ISCs are essential for formation of a functional epithelial barrier, and disruptions to the epithelium often lead to severe inflammatory disease.

Flies are an ideal animal model to scrutinize links between stem cell immunity and gut barrier organization^49–52^. Antibacterial immunity in the *Drosophila* gut includes prominent inputs from the IMD pathway, a germline-encoded defense with striking similarities to the vertebrate antibacterial NOD2 pathway. Like vertebrates, the fly midgut is maintained by multipotent progenitors that produce mature enterocyte and endocrine lineages in response to signals from Notch, EGF, and BMP ligands. Notably, transcriptomic and physiological studies point to elaborate connections between intestinal IMD pathway activity and cellular functions in the fly^32,36,53–60^. Thus, flies are an obvious genetic tool for foundational discoveries about the impacts of critical immune pathways on intestinal epithelial dynamics.

We used the midgut as a model system to determine how progenitor cell-specific Rel activity maintains the epithelial barrier. Our initial experiments agree with prior studies that implicated progenitor cell immunity in age-dependent decay of epithelial organization^58^. However, our work went further by characterizing effects of progenitor cell Rel on epithelial regeneration after challenges with DSS, an injurious, colitogenic polysaccharide that damages the epithelium and prompts a rapid regenerative response in the gut. We found that, alongside EGF, JAK-STAT and Notch signals, DSS also induced immune responses that included progenitor cell-specific expression of Rel signature genes such as *pirk*, *PGRP-SD*, and *PGRP*-*LB*, indicating induction of immune signals during enteric stress. ISC-specific loss of *Rel* led to stress-dependent progenitor death that impaired damage-responsive generation of pre-enterocytes, and enhanced fly sensitivity to DSS. Our observations implicate IMD in the survival of DSS-exposed stem cells and raises questions that we feel merit consideration in future studies.

How do progenitors detect IMD ligands during periods of stress? Typically, Rel responds to bacterial DAP-type peptidoglycan via transmembrane and intracellular PGRPs^61–68^, although commensal-derived acetate also activates Rel in midgut enteroendocrine cells^69^. A dense, chitinous peritrophic matrix combined with a contiguous layer of mature epithelial cells protects ISCs from direct contact with the lumen, while basement membrane and visceral muscle prevent extensive environmental interactions with the ISC basolateral surface^70^. As a result, it is unclear how luminal DSS prompts Rel activation in such a shielded cell. In a simple model, DSS may breach the barrier to the extent that gut-resident bacterial patterns access immune receptors on progenitors. If paracellular leakage of bacterial patterns initiates immune responses in progenitors, it would be of interest to test how defined bacterial communities, including absence of gut bacteria, modify stem cell survival and barrier regeneration after a DSS challenge. In this context, we note that researchers have identified enteric pathogens that differentially regulate epithelial regeneration and IMD signals. With a simple, manipulable microbiome, and straightforward infection protocols, flies are an excellent model to ask how hosts, commensals, and pathogens interact to shape stress-dependent epithelial regeneration^10,20,56,71^.

How does Rel modify stem cell survival? Loss of Rel enhances stem cell death and increases expression of apoptotic modifiers after exposure to DSS, while inhibition of apoptosis restores regenerative capacity to Rel-deficient progenitors, indicating a requirement for immune signals in stem cell survival during DSS exposure. Our work overlaps with literature that implicates immune signals in cell viability, but the precise mechanism by which Rel modifies epithelial survival requires clarification. As Rel has broad effects on gene expression within the midgut, including genes linked to stress responses and development^32^, it is possible that the transcriptional activity of Rel directly modifies expression of genes involved in cell survival. However, we cannot exclude the possibility that Rel controls survival via intermediaries. For example, our transcriptional data point to unanticipated effects of Rel on pathways linked to regeneration: inactivation of Rel suppresses DSS-dependent EGF and Notch signals without notable effects on JAK-STAT activity. Moving forward, it will be of interest to test if interactions between Rel and EGF, Notch or JAK-STAT are important for DSS-dependent control of stem cell viability. Beyond, EGF, Notch, and JAK-STAT, we feel it will be pertinent to test the tumor suppressor BMP pathway as a potential mediator of Rel-dependent stem cell survival. BMP regulates epithelial patterning^28,38,72–77^; DSS enhances expression of the BMP pathway elements *Mad* and *brk* in progenitors (Figure 3D); and *Vibrio cholerae* engages a Rel-BMP axis that blocks ISC proliferation^78^, suggesting possible interactions between Rel and BMP in the regulation of cell viability after exposure to DSS.

What about enteroendocrine cells? Our work focussed on stress-dependent epithelial regeneration, a process dominated by production of large, polyploid enterocytes that constitute the bulk of adult midgut. However, enteroendocrine cells are essential elements of a mature intestine, with established roles in detection of bacterial metabolites, and regulation of stem cell dynamics^79^. In flies, enteroendocrine cells constitute roughly 10% of the midgut epithelium and secrete peptide hormones that modify gut behaviour locally, but also organ function at distant sites^80^. Luminal factors that include microbial products shape the identity, number, and functional state of enteroendocrine cells, and recent studies implicate progenitor cell-restricted Rel activity in enteroendocrine cell development^36,53^. As acute bacterial infections have long-term consequences for the specification of enteroendocrine subtypes^81^, we believe it will be of interest to determine what effects stem cell-specific Rel activity has on enteroendocrine cell generation after a challenge, and whether any effects are transitory or permanent in nature.

Finally, we note that our work focussed on the adult female midgut, a common practice within the *Drosophila* community. As there is an abundance of evidence the gut structure, function, immunity and proliferation are sexually dimorphic^82,83,84,85^, we believe it would be of interest to test effects of Rel inactivation on the male ISC response to extrinsic challenges.

Together our observations suggest that Relish acts in ISCs to control cell survival in the face of an extrinsic insults, and that Rel-dependent ISC survival is essential for effective epithelial repair. Given the constant environmental fluctuations the intestinal lumen is exposed to, we suggest that immune signaling in ISCs is used as an adaptive mechanism to tune cell survival, differentiation and proliferation to the specific needs of the epithelium.

## MATERIALS AND METHODS

### Drosophila husbandry

*Drosophila* crosses were setup and maintained at 18°C on standard corn meal food (Nutri-Fly Bloomington formulation; Genesse Scientific). All experimental flies were virgin female and kept at a 12h:12h light dark cycle throughout. Upon eclosion, flies were kept at 18°C then shifted to the appropriate temperature once 25-30 flies per vial was obtained. Fly lines used in this study were: *w ; esg-GAL4,tubGAL80ts,UAS-GFP* (*esg^ts^)* (Bruce Edgar), *w ; esg-GAL4,UAS-2xEYFP;Su(H)GBE-GAL80,tub-GAL80^ts^* (*ISC^ts^*) (Bruce Edgar), *w ; esg-GAL4, tub-GAL80^ts^, UAS-GFP* ; *UAS-flp, Act>CD2>GAL4* (*esg F/O*) (Bruce Edgar), *w ; esg-GAL4,UAS-CFP, Su(H)-GFP;tubGal80^ts^ (esgts,UAS-CFP,SuH-GFP)* (Lucy O’Brien), *w^1118^* (VDRC #60000), *Rel^RNAi^* (VDRC #49413), *key^RNAi^* (VDRC #7723).

### Immunofluorescence

Intestines were dissected in PBS, fixed in 8% formaldehyde for 20 minutes, washed in PBS 0.2% Triton-X (PBST) then blocked in PBST with 3% BSA for 1hr at room temperature. Primary antibodies were incubated in PBST with BSA overnight at 4°C. The following day guts were washed in PBST then secondary antibody incubations were done in conjunction with DNA stain for 1 hour at room temperature in PBST with BSA, washed with PBST then again with PBS. Primary antibodies used: anti-prospero (1/100; Developmental Studies Hybridoma Bank (DSHB) MR1A), anti-armadillo (1/100;DSHB N2 7A1), chicken anti-GFP (1/2000; Invitrogen PA1-9533), anti-phospho-histone3 (1/1000; Millipore 06-570), anti-Delta (1/100; DSHB C594.9B). Secondary antibodies used: goat anti-chicken 488 (1/1000; Invitrogen A11039), goat anti-mouse 568 (1/1000; Invitrogen A11004), goat anti-rabbit 568 (1/1000; Invitrogen A11011), goat anti-mouse 647 (1/1000; Invitrogen A21235), and goat anti-rabbit 647 (1/1000; Invitrogen A21244). DNA stains used: Hoechst (1/1000; Molecular Probes H-3569). Apoptotic cells were detected in dissected guts using the TMR red In Situ Cell Death Detection Kit (Roche; 12156792910) following standard kit staining protocol. Briefly, guts were washed in PBS following secondary antibody then stained with 100μL of TUNEL solution for 1hr at 37°C then washed twice with PBS. Intestines were mounted on slides using Fluoromount (Sigma; F4680). For every experiment, images were obtained of the posterior midgut region (R4/5) of the intestine with a spinning disk confocal microscope (Quorum WaveFX). PH3+ cells were counted through the entire midgut.

### DSS treatment

DSS was prepared by dissolving DSS (Sigma 42867) in a PBS 5% sucrose solution, filter sterilized and kept in the freezer for up to two weeks. Flies were raised on a 5% DSS solution unless stated otherwise. DSS vials were prepped by covering normal fly food with circular filter paper (Whatman, Grade 3, 23mm, 1003-323) and adding 150μL of DSS, or control (PBS 5% sucrose) solutions. Flies were flipped daily onto fresh DSS, or control solution for 48hr for all experiments except for DSS survival which was over the course of 10 days.

### Lifespan

For longevity, 30 virgin females per vial were raised at 29°C and dead flies were counted every 1-3 days and vials were flipped 3 times per week to fresh standard food. For DSS survival experiments flies were placed on fresh 10% DSS daily for the course of the survival experiment and deaths were counted daily.

### Data visualization and Statistical analysis

Figures were constructed using R (version 4.1.2) via R studio with easyggplot2 (version 1.0.0.9000) or ggplot2 (version 3.3.5). Statistical analysis was performed in R. Figures were assembled in Adobe Illustrator.

### Sample prep for single cell RNA sequencing

Preparation of single-cell intestinal suspension was made following previous methods (31, 44). Flies were raised for 10 days at 29°C then treated with 3% DSS or sucrose/PBS solution for 48hrs. Batches of five *Drosophila* midguts were dissected at once then transferred to 1% BSA in DEPC treated PBS. Once 30 midguts were obtained for each condition they were transferred to a 1.5mL tube with 200μL of DEPC/PBS with 1mg/mL Elastase (Sigma, E0258) and chopped into pieces with small dissecting scissors. After mechanical disruption, tubes were incubated at 27°C for 40min with gentle pipetting every 10min. 22μL of 10%BSA in DEPC/PBS solution was added to stop the enzymatic disruption then cells were pelleted by spinning at 300g for 15min at 4°C. Cell pellet was resuspended in 200μL of 0.04% BSA in DEPC/PBS then filtered through a 70μm filter. Live cells were enriched using OptiPrep Density Gradient Medium (Sigma, D1556). Filtered cells were mixed with 444μL of 40% iodixanol (2:1 OptiPrep:0.04% BSA DEPC/PBS) then transferred to a 15mL tube. Another 5.36mL of 40% iodixanol was added and mixed. A 3mL layer of 22% iodixanol was added on top then an additional 0.5mL layer of 0.04% BAS in PBS/DEPC was added. Tubes were spun at 800g for 20min at 20°C then the top interface containing live cells (∼500μL) was collected. Live cells were diluted with 1mL of 0.04% BSA in DEPC/PBS. Remaining iodixanol was removed by pelleting cells at 300g for 10min at 4?C and removing supernatant. Cell pellet was resuspended in remaining 0.04% BSA DEPC/PBS solution (∼40μL) and cell counts and viability was determined using a hemocytometer. Cell viability was as follows: *ISCts/+* unchallenged = 96%, *ISCts/+* DSS = 95%, *ISCts/relRNAi* unchallenged = 97%, *ISCts/relRNAi* DSS = 86%. Libraries were generated using 10X Genomics Single-cell Transcriptome Library kit and sent to Novogene for sequencing.

### Bioinformatics

Raw sequencing data from Novogene was aligned to the *Drosophila* reference transcriptome (FlyBase, r6.30) using Cell Ranger v3.0 with the EYFP sequence appended to generate feature-barcode matrices. The resulting matrices were analyzed using Seurat (v4.1.0)(45, 46) in R. Cells with <200 or >3500 features and cells with >20% mitochondrial reads were removed to reduce number of low quality cells or doublets. Expression values were normalized and data clustering was performed at a resolution of 0.4 with 30 principal components. Clusters were identified using established markers and previous *Drosophila* intestine single-cell analysis (www.flyrnai.org/scRNA). GO term analysis was performed using DAVID.

### Data Availability

Gene expression data is deposited on NCBI under the accession number PRJNA873108.

---

**Supplementary Figure 1.**
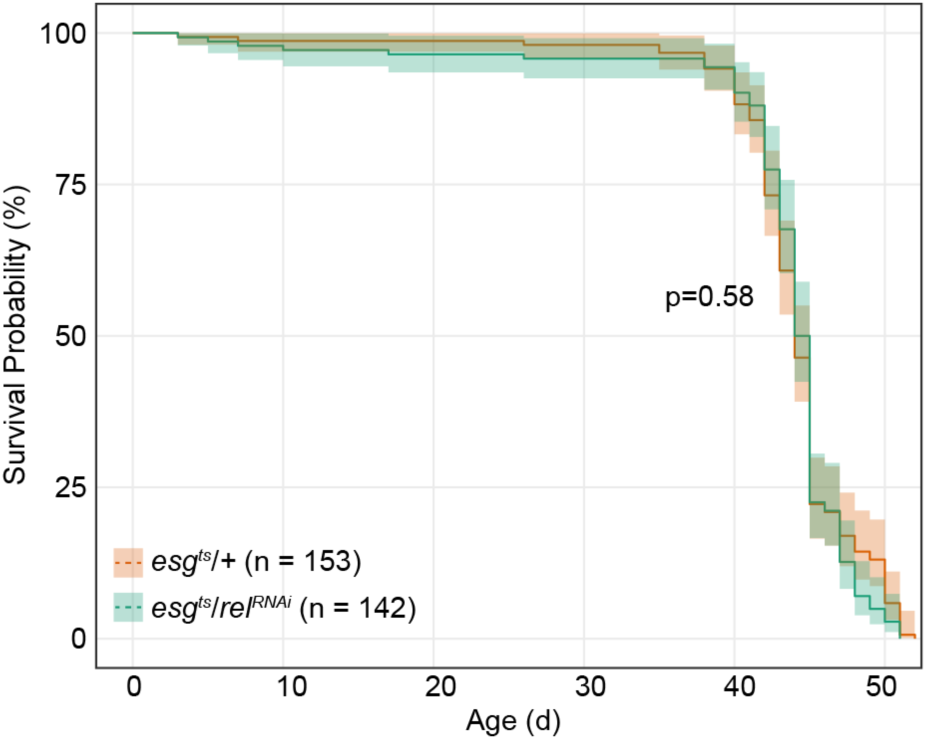
Lifespan analysis of *esg^ts^/+* and *esg^ts^/Rel^RNAi^* adult female flies raised under conventional laboratory conditions. Significance calculated using log rank test.

**Supplementary Figure 2.**
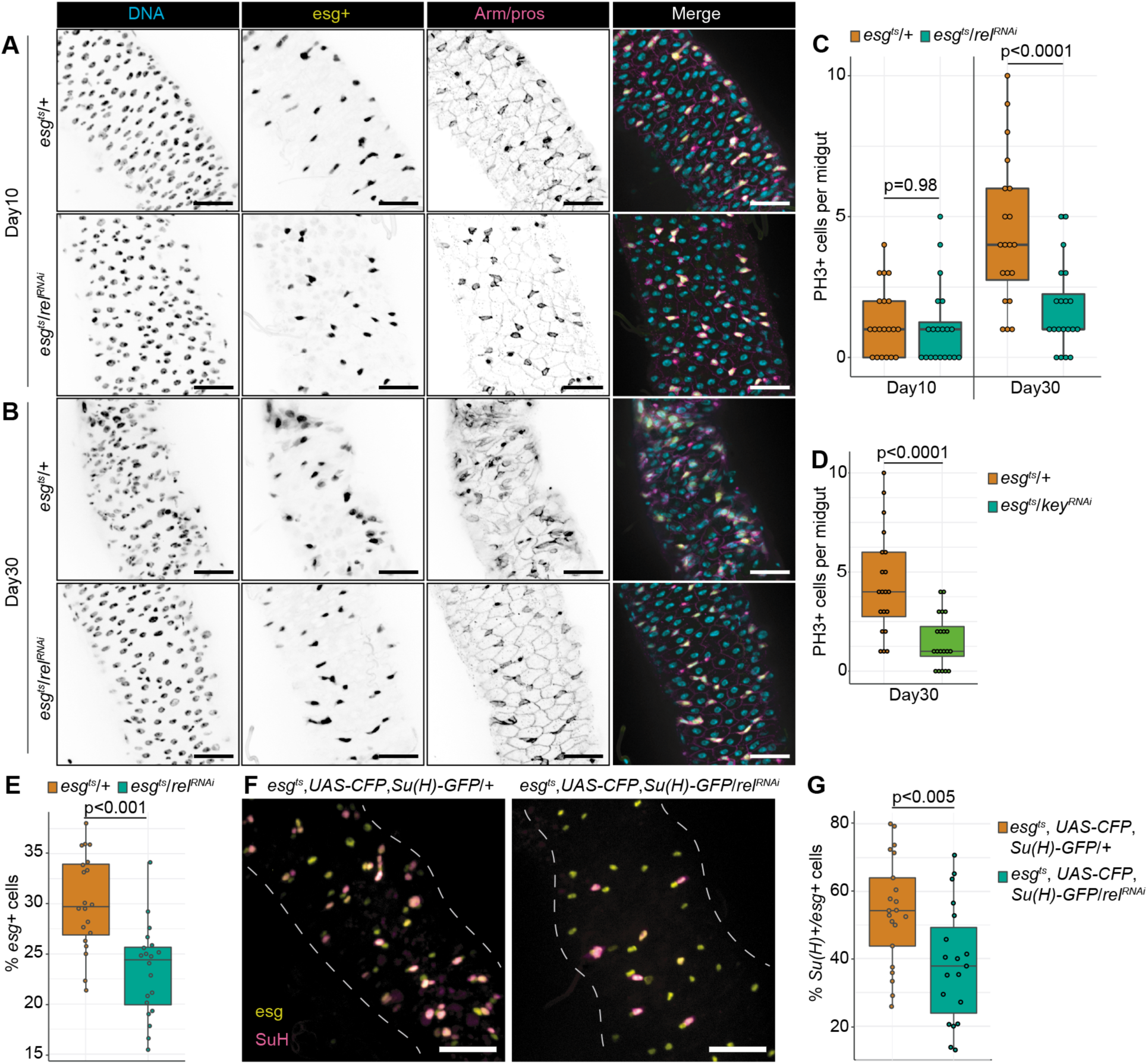
**NF-**κ**B regulates intestinal stem cell proliferation. (A)** Posterior midgut of flies after 10 days of progenitor-specific NF-κB depletion (*esg^ts^/Rel^RNAi^*) compared to wild-type (*esg^ts^/+).* DNA labelled with Hoechst (cyan), progenitors labelled with esg (yellow), enteroendocrine cells labeled by pros (magenta) and cell borders labelled by Armadillo (magenta). **(B)** Posterior midgut of flies after 30 days of progenitor-specific *Rel* depletion compared to wild-type. **(C)** PH3+ mitotic cells per midgut of wild-type and progenitor-specific *Rel* depleted flies after 10 and 30 days. Significance calculated using ANOVA followed by pairwise Tukey tests. **(D)** PH3+ cells per midgut after progenitor-specific knockdown of IKKγ homolog *kenny* (*key*) **(E)** Proportion of esg+ progenitors per nucleus in 30-day old flies upon progenitor-specific *Rel* knockdown. **(F)** Images from *esg^ts^, UAS-CFP, Su(H)-GFP* flies after 30 days of progenitor-specific *Rel* depletion. Progenitors labelled with esg (yellow) and Notch positive enterocyte precursors labelled with Su(H) (magenta). **(G)** Proportion of Su(H)+ enterocyte precursors within the progenitor pool upon progenitor-specific NFκB knockdown. For D, E and G significance found using Student’s t test. Scale bars for A and B are 25μm.

**Supplementary Figure 3.**
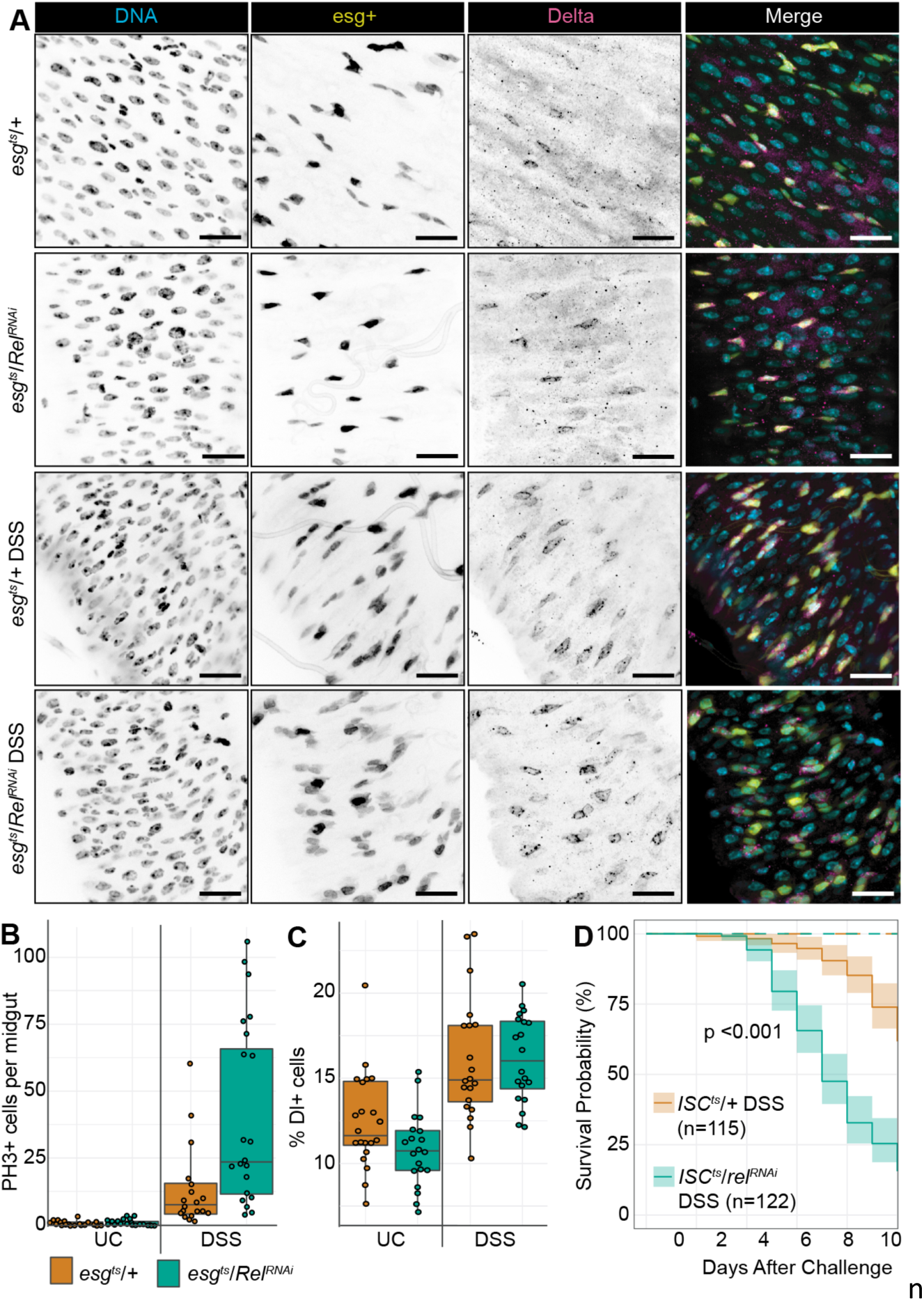
DSS Damages the Intestinal Epithelium and Induces a Proliferative Response in Progenitor Cells. **(A)** Posterior midgut of 10 days old unchallenged (UC) *esg^ts^/+* flies fed PBS/5% sucrose solution for 48hrs; and DSS-treated *esg^ts^/+* or *esg^ts^/rel^RNAi^* flies fed a 3% DSS solution in 5% sucrose for 48hrs. Scale bars = 25μm. DNA labelled by Hoechst (cyan), esg+ progenitors (yellow), ISCs labelled with Delta (magenta). **(B)** PH3+ cells per midgut of 48hr UC and DSS treated *esg^ts^/+* flies. **(C)** Percentage Delta (Dl)-positive cells per midgut of unchallenged (UC) *esg^ts^/+* or *esg^ts^/rel^RNAi^* flies fed a sucrose solution or a DSS solution for 48h. **(D)** Survival of *ISC^ts^/+* and *ISC^ts^/Rel^RNAi^* flies treated with DSS. Significance calculated with a log rank test.

**Supplementary Figure 4.**
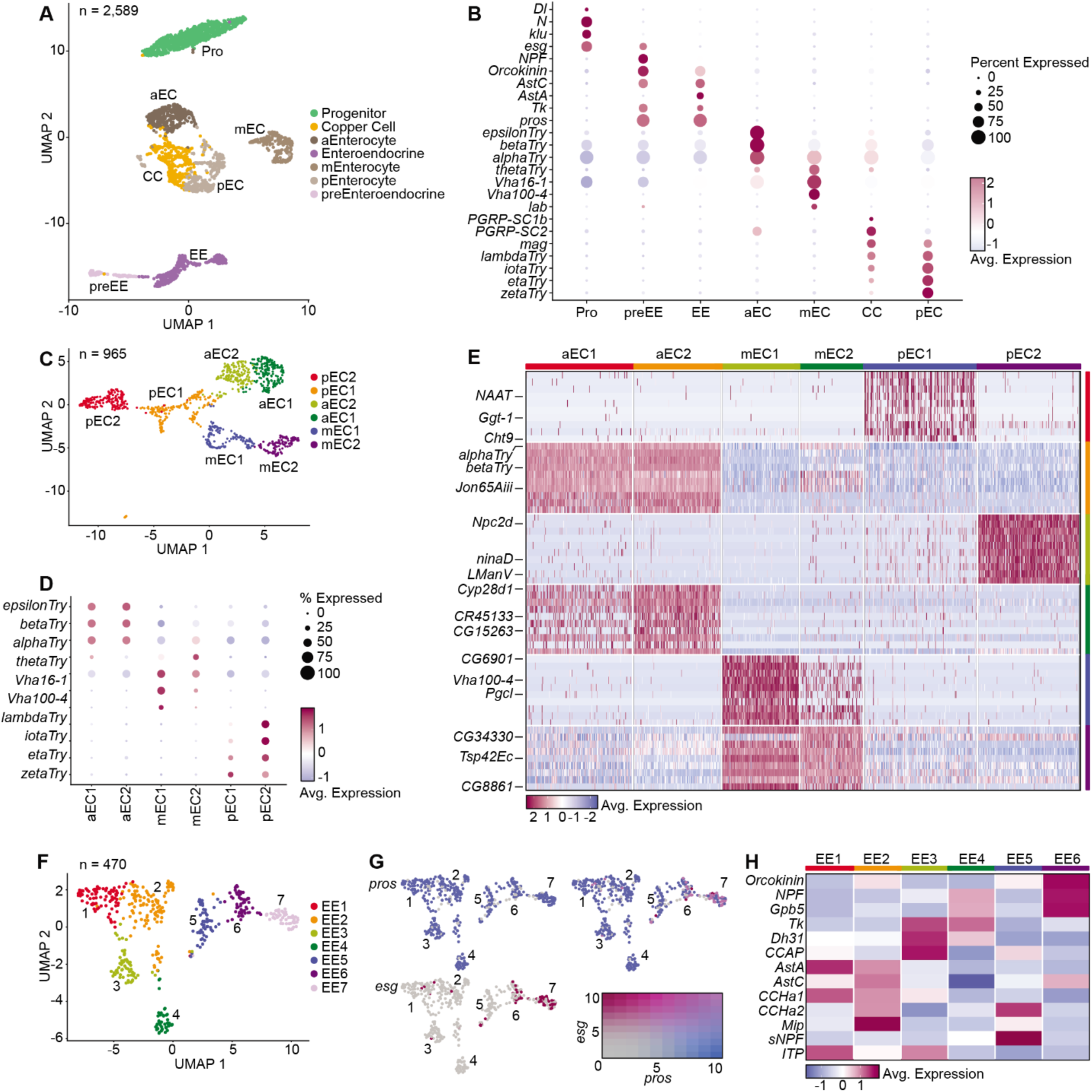
Single Cell Gene Expression Atlas of the Adult *Drosophila* Midgut. **(A)** UMAP representation of cell types found in the intestines of 10-day-old *ISC^ts/+^* flies. **(B)** Dotplot representation of the expression levels of established markers in the indicated cell types. **(C)** UMAP clustering of six enterocyte subtypes in the adult midgut **(D)** Dotplot showing the relative expression of known anterior (aEC), middle (mEC), and posterior (pEC) enterocyte markers in each cluster. **(E)** Heatmap of marker gene expression for each cluster, showing prominent cell cluster markers. **(F-G)** UMAP clustering of seven prospero-positive enteroendocrine subtypes in the adult midgut **(F)**, with a feature plot **(G)** showing enriched expression of the progenitor cell marker esg in cluster seven. **(H)** Heatmap showing relative expression of enteroendocrine-derived peptide hormones in each mature enteroendocrine cell subset. For each graph, pro = progenitor; preEE = pre-enteroendocrine; ee = enteroendocrine; aEC, mEC, and pEC = anterior, middle and posterior enterocytes, respectively; and CC = copper cells.

**Supplementary Figure 5.**
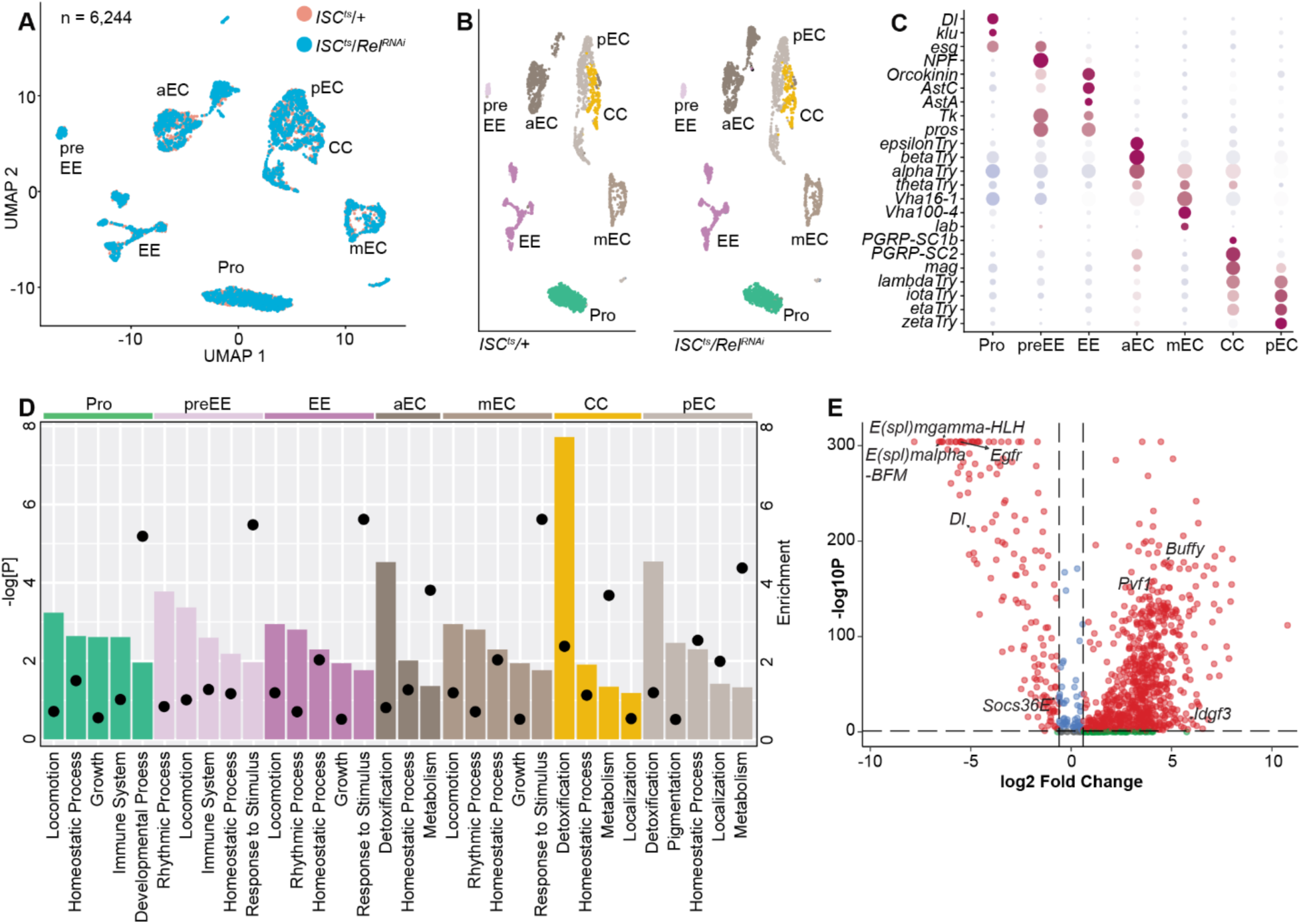
Inactivation of Rel in Stem Cells Impacts Transcriptional States Throughout the Gut. (A-B) Integrated (A), and individual (B) UMAP projections of cell types found in midguts from ten-day-old *ISC^ts/+^* and *ISC^ts^/Rel^RNAi^* flies. **(C)** Dotplot representation of the expression levels of established markers in the indicated cell types. **(D)** Representation of the top five downregulated gene ontology terms in *ISC^ts^/Rel^RNAi^* midguts relative to ISCts/+ controls for each cell type. Relative enrichment score for each term is indicated by bar length and the negative log p-values are shown with closed circles. **(E)** Volcano plot showing progenitor-specific gene expression in *ISC^ts^/Rel^RNAi^* midguts relative to *ISC^ts/+^* controls. A set of significantly modified genes are indicated. For each graph, pro = progenitor; preEE = pre-enteroendocrine; ee = enteroendocrine; aEC, mEC, and pEC = anterior, middle and posterior enterocytes, respectively; and CC = copper cells.

**Supplementary Figure 6.**
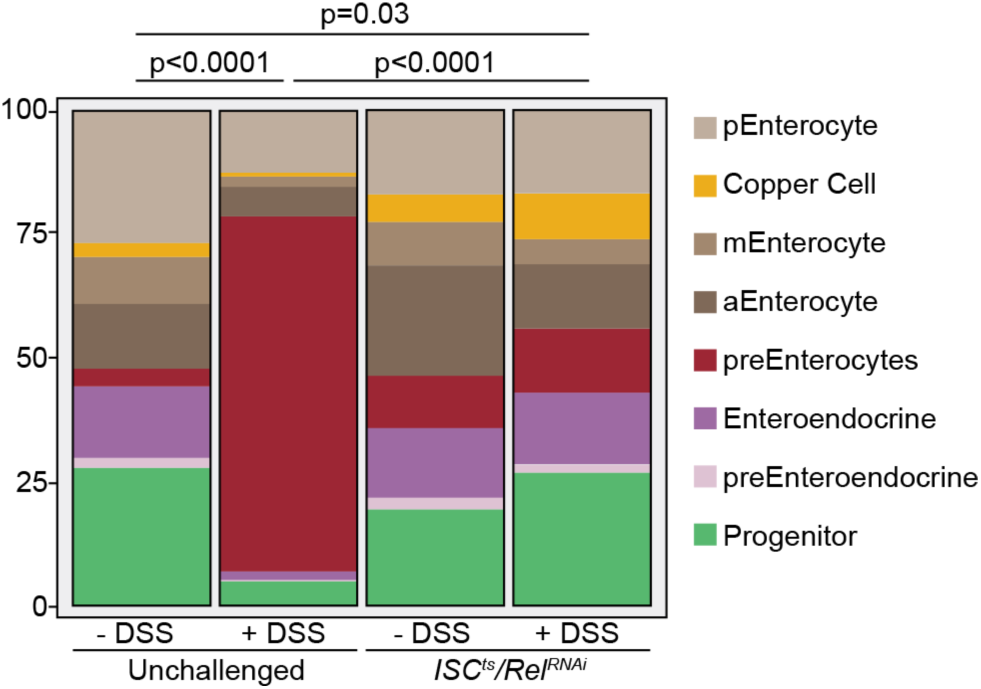
Cluster type abundance from UMAPs seen in Supplementary Figures 4-5, and Figure 2. Significance for the preEC proportion was calculated using a chi squared test followed by proportions test.

## CITATIONS

1. Ferguson, M. & Foley, E. Microbial recognition regulates intestinal epithelial growth in homeostasis and disease. FEBS 1–26 (2021) doi:10.1111/febs.15910.

2. Ullman, T. A. & Itzkowitz, S. H. Intestinal inflammation and cancer. Gastroenterology 140, 1807–1816 (2011).

3. Hugot, J. P. et al. Mapping of a susceptibility locus for Crohn’s disease on chromosome 16. Nature 379, 821–823 (1996).

4. Ogura, Y. et al. A frameshift mutation in NOD2 associated with susceptibility to Crohn’s disease. Nature 411, 603–606 (2001).

5. Jostins, L. et al. Host-microbe interactions have shaped the genetic architecture of inflammatory bowel disease. Nature 491, 119–124 (2012).

6. Hugot, J. P. et al. Association of NOD2 leucine-rich repeat variants with susceptibility to Crohn’s disease. Nature 411, 599–603 (2001).

7. Couturier-Maillard, A. et al. NOD2-mediated dysbiosis predisposes mice to transmissible colitis and colorectal cancer. J. Clin. Invest. 123, 700–711 (2013).

8. Choi, P. M. & Zelig, M. P. Similarity of colorectal cancer in Crohn’s disease and ulcerative colitis: implications for carcinogenesis and prevention. Gut 35, 950–954 (1994).

9. Miguel-Aliaga, I., Jasper, H. & Lemaitre, B. Anatomy and Physiology of the Digestive Tract of Drosophila melanogaster. Genetics 210, 357–396 (2018).

10. Buchon, N., Broderick, N. A. & Lemaitre, B. Gut homeostasis in a microbial world: insights from Drosophila melanogaster. Nat. Rev. Microbiol. 11, 615–626 (2013).

11. Ludington, W. B. & Ja, W. W. Drosophila as a model for the gut microbiome. PLOS Pathog. 16, e1008398 (2020).

12. Micchelli, C. A. & Perrimon, N. Evidence that stem cells reside in the adult Drosophila midgut epithelium. Nature 439, 475–479 (2006).

13. Ohlstein, B. & Spradling, A. The adult Drosophila posterior midgut is maintained by pluripotent stem cells. Nature 439, 470–474 (2006).

14. O’Brien, L. E., Soliman, S. S., Li, X. & Bilder, D. Altered Modes of Stem Cell Division Drive Adaptive Intestinal Growth. Cell 147, 603–614 (2011).

15. Goulas, S., Conder, R. & Knoblich, J. A. The Par complex and integrins direct asymmetric cell division in adult intestinal stem cells. Cell Stem Cell 11, 529–540 (2012).

16. Perdigoto, C. N., Schweisguth, F. & Bardin, A. J. Distinct levels of Notch activity for commitment and terminal differentiation of stem cells in the adult fly intestine. Dev. Camb. Engl. 138, 4585–4595 (2011).

17. Joly, A. & Rousset, R. Tissue Adaptation to Environmental Cues by Symmetric and Asymmetric Division Modes of Intestinal Stem Cells. Int. J. Mol. Sci. 21, 6362 (2020).

18. Ohlstein, B. & Spradling, A. Multipotent Drosophila Intestinal Stem Cells Specify Daughter Cell Fates by Differential Notch Signaling. Science 315, 988–993 (2007).

19. Ngo, S., Liang, J., Su, Y.-H. & O’Brien, L. E. Disruption of EGF Feedback by Intestinal Tumors and Neighboring Cells in Drosophila. Curr. Biol. CB 30, 1537–1546.e3 (2020).

20. Buchon, N., Broderick, N. A., Chakrabarti, S. & Lemaitre, B. Invasive and indigenous microbiota impact intestinal stem cell activity through multiple pathways in Drosophila. Genes Dev. 23, 2333–2344 (2009).

21. Bardin, A. J., Perdigoto, C. N., Southall, T. D., Brand, A. H. & Schweisguth, F. Transcriptional control of stem cell maintenance in the Drosophila intestine. Dev. Camb. Engl. 137, 705–714 (2010).

22. Sallé, J. et al. Intrinsic regulation of enteroendocrine fate by Numb. EMBO J. 36, 1928–1945 (2017).

23. Cordero, J. B., Stefanatos, R. K., Myant, K., Vidal, M. & Sansom, O. J. Non-autonomous crosstalk between the Jak/Stat and Egfr pathways mediates Apc1-driven intestinal stem cell hyperplasia in the Drosophila adult midgut. Dev. Camb. Engl. 139, 4524–4535 (2012).

24. Nászai, M. et al. RAL GTPases mediate EGFR-driven intestinal stem cell proliferation and tumourigenesis. eLife 10, e63807 (2021).

25. Jin, Y. et al. EGFR/Ras Signaling Controls Drosophila Intestinal Stem Cell Proliferation via Capicua-Regulated Genes. PLoS Genet. 11, 1–27 (2015).

26. Biteau, B. & Jasper, H. EGF signaling regulates the proliferation of intestinal stem cells in Drosophila. Development 138, 1045–1055 (2011).

27. Jiang, H. et al. Cytokine/Jak/Stat signaling mediates regeneration and homeostasis in the Drosophila midgut. Cell 137, 650–657 (2009).

28. Tian, A. & Jiang, J. Intestinal epithelium-derived BMP controls stem cell self-renewal in Drosophila adult midgut. eLife 3, 1–23 (2014).

29. Guo, Z., Driver, I. & Ohlstein, B. Injury-induced BMP signaling negatively regulates Drosophila midgut homeostasis. JCB 201, 945–961 (2013).

30. Takashima, S. et al. Development of the Drosophila entero-endocrine lineage and its specification by the Notch signaling pathway. Dev. Biol. 353, 161–172 (2011).

31. Erkosar, B. et al. Drosophila microbiota modulates host metabolic gene expression via IMD/NF-κB signaling. PloS One **G**, e94729 (2014).

32. Broderick, N. A., Buchon, N. & Lemaitre, B. Microbiota-Induced Changes in Drosophila melanogaster Host Gene Expression and Gut Morphology. mBio 5, 1–13 (2014).

33. Buchon, N., Broderick, N. A., Poidevin, M., Pradervand, S. & Lemaitre, B. Drosophila Intestinal Response to Bacterial Infection: Activation of Host Defense and Stem Cell Proliferation. Cell Host Microbe 5, 200–211 (2009).

34. Myllymäki, H., Valanne, S. & Rämet, M. The Drosophila imd signaling pathway. J. Immunol.192, 3455–62 (2014).

35. Shin, M. et al. Immune regulation of intestinal-stem-cell function in Drosophila. Stem Cell Rep. 17, 741–755 (2022).

36. Liu, X. et al. Microbes affect gut epithelial cell composition through immune-dependent regulation of intestinal stem cell differentiation. Cell Rep. 38, 110572 (2022).

37. Hung, R.-J. et al. A cell atlas of the adult Drosophila midgut. Proc. Natl. Acad. Sci. 117, 1514–1523 (2020).

38. Boutros, M., Agaisse, H. & Perrimon, N. Sequential Activation of Signaling Pathways during Innate Immune Responses in Drosophila. Dev. Cell 3, 711–722 (2002).

39. Vandehoef, C., Molaei, M. & Karpac, J. Dietary Adaptation of Microbiota in *Drosophila* Requires NF-κB-Dependent Control of the Translational Regulator 4E-BP. Cell Rep. 31, 107736 (2020).

40. Rodriguez-Fernandez, I. A., Tauc, H. M. & Jasper, H. Hallmarks of aging Drosophila intestinal stem cells. Mech. Ageing Dev. 190, 111285 (2020).

41. Martin, J. L. et al. Long-term live imaging of the Drosophila adult midgut reveals real-time dynamics of division, differentiation and loss. eLife 7, e36248 (2018).

42. Amcheslavsky, A., Jiang, J. & Ip, Y. T. Tissue Damage-Induced Intestinal Stem Cell Division in Drosophila. Cell Stem Cell 4, 49–61 (2009).

43. Ren, F. et al. Hippo signaling regulates Drosophila intestine stem cell proliferation through multiple pathways. Proc. Natl. Acad. Sci. U. S. A. 107, 21064–21069 (2010).

44. Naik, S., Larsen, S. B., Cowley, C. J. C, Fuchs, E. Two to Tango: Dialog between Immunity and Stem Cells in Health and Disease. Cell 175, 908–920 (2018).

45. Neal, M. D. et al. Toll-like Receptor 4 Is Expressed on Intestinal Stem Cells and Regulates Their Proliferation and Apoptosis via the p53 Up-regulated Modulator of Apoptosis. J. Biol. Chem. 287, 37296–37308 (2012).

46. Du, J. et al. Zymosan-A promotes the regeneration of intestinal stem cells by upregulating ASCL2. Cell Death Dis. 13, 1–11 (2022).

47. Afrazi, A. et al. Toll-like Receptor 4-mediated Endoplasmic Reticulum Stress in Intestinal Crypts Induces Necrotizing Enterocolitis *. J. Biol. Chem. 289, 9584–9599 (2014).

48. Hageman, J. H. et al. Intestinal Regeneration: Regulation by the Microenvironment. Dev. Cell 54, 435–446 (2020).

49. Khan, S. A., Kojour, M. A. M. C, Han, Y. S. Recent trends in insect gut immunity. Front. Immunol.14, (2023).

50. Tafesh-Edwards, G. C, Eleftherianos, I. The role of Drosophila microbiota in gut homeostasis and immunity. Gut Microbes 15, 2208503 (2023).

51. Buchon, N., Silverman, N. C, Cherry, S. Immunity in Drosophila melanogaster — from microbial recognition to whole-organism physiology. Nat. Rev. Immunol. 14, 796–810 (2014).

52. Capo, F., Wilson, A. C, Di Cara, F. The Intestine of Drosophila melanogaster: An Emerging Versatile Model System to Study Intestinal Epithelial Homeostasis and Host-Microbial Interactions in Humans. Microorganisms 7, 336 (2019).

53. Shin, M. et al. Immune regulation of intestinal-stem-cell function in Drosophila. Stem Cell Rep. 17, 741–755 (2022).

54. Zhai, Z., Boquete, J.-P. C, Lemaitre, B. Cell-Specific Imd-NF-κB Responses Enable Simultaneous Antibacterial Immunity and Intestinal Epithelial Cell Shedding upon Bacterial Infection. Immunity 48, 897–910.e7 (2018).

55. Broderick, N. A. Friend, foe or food? Recognition and the role of antimicrobial peptides in gut immunity and Drosophila–microbe interactions. Philos. Trans. R. Soc. B Biol. Sci. 371, 20150295 (2016).

56. Lesperance, D. N. & Broderick, N. A. Microbiomes as modulators of Drosophila melanogaster homeostasis and disease. Curr. Opin. Insect Sci. **39**, 84–90 (2020).

57. Barretto, E. C., Polan, D. M., Beevor-Potts, A. N., Lee, B. C, Grewal, S. S. Tolerance to Hypoxia Is Promoted by FOXO Regulation of the Innate Immunity Transcription Factor NF-κB/Relish in Drosophila. Genetics 215, 1013–1025 (2020).

58. Guo, L., Karpac, J., Tran, S. L. C, Jasper, H. PGRP-SC2 Promotes Gut Immune Homeostasis to Limit Commensal Dysbiosis and Extend Lifespan. Cell 156, 109–122 (2014).

59. Yamashita, K. et al. Activation of innate immunity during development induces unresolved dysbiotic inflammatory gut and shortens lifespan. Dis. Model. Mech. 14, dmm049103 (2021).

60. Kosakamoto, H. et al. Local Necrotic Cells Trigger Systemic Immune Activation via Gut Microbiome Dysbiosis in Drosophila. Cell Rep. 32, (2020).

61. Kaneko, T. et al. PGRP-LC and PGRP-LE have essential yet distinct functions in the drosophila immune response to monomeric DAP-type peptidoglycan. Nat. Immunol. 7, 715–723 (2006).

62. Kaneko, T. et al. Monomeric and polymeric gram-negative peptidoglycan but not purified LPS stimulate the Drosophila IMD pathway. Immunity 20, 637–649 (2004).

63. Leulier, F. et al. The Drosophila immune system detects bacteria through specific peptidoglycan recognition. Nat. Immunol. 4, 478–484 (2003).

64. Takehana, A. et al. Peptidoglycan recognition protein (PGRP)-LE and PGRP-LC act synergistically in Drosophila immunity. EMBO J. 23, 4690–4700 (2004).

65. Gottar, M. et al. The Drosophila immune response against Gram-negative bacteria is mediated by a peptidoglycan recognition protein. Nature 416, 640–644 (2002).

66. Choe, K.-M., Werner, T., Stöven, S., Hultmark, D. C, Anderson, K. V. Requirement for a peptidoglycan recognition protein (PGRP) in Relish activation and antibacterial immune responses in Drosophila. Science 296, 359–362 (2002).

67. Rämet, M., Manfruelli, P., Pearson, A., Mathey-Prevot, B. C, Ezekowitz, R. A. B. Functional genomic analysis of phagocytosis and identification of a Drosophila receptor for E. coli. Nature 416, 644–648 (2002).

68. Choe, K.-M., Lee, H. C, Anderson, K. V. Drosophila peptidoglycan recognition protein LC (PGRP-LC) acts as a signal-transducing innate immune receptor. Proc. Natl. Acad. Sci. 102, 1122–1126 (2005).

69. Jugder, B.-E., Kamareddine, L. C, Watnick, P. I. Microbiota-derived acetate activates intestinal innate immunity via the Tip60 histone acetyltransferase complex. Immunity 54, 1683–1697.e3 (2021).

70. Galenza, A. et al. Basal stem cell progeny establish their apical surface in a junctional niche during turnover of an adult barrier epithelium. Nat. Cell Biol. 25, 658–671 (2023).

71. Barron, A. J. et al. Microbiome-derived acidity protects against microbial invasion in Drosophila. Cell Rep. 43, 114087 (2024).

72. Tracy Cai, X., et al. AWD regulates timed activation of BMP signaling in intestinal stem cells to maintain tissue homeostasis. Nat. Commun. 10, 2988 (2019).

73. Tian, A., Wang, B. C, Jiang, J. Injury-stimulated and self-restrained BMP signaling dynamically regulates stem cell pool size during Drosophila midgut regeneration. Proc. Natl. Acad. Sci. 114, E2699–E2708 (2017).

74. Guo, Z., Driver, I. C, Ohlstein, B. Injury-induced BMP signaling negatively regulates Drosophila midgut homeostasis. J. Cell Biol. 201, 945–961 (2013).

75. Driver, I. & Ohlstein, B. Specification of regional intestinal stem cell identity during Drosophila metamorphosis. Development 141, 1848–1856 (2014).

76. Zhou, J. et al. Dpp/Gbb signaling is required for normal intestinal regeneration during infection. Dev. Biol. 3GG, 189–203 (2015).

77. Christensen, C. F., Laurichesse, Q., Loudhaief, R., Colombani, J. C, Andersen, D. S. Drosophila activins adapt gut size to food intake and promote regenerative growth. Nat. Commun. 15, 273 (2024).

78. Xu, X. & Foley, E. Vibrio cholerae arrests intestinal epithelial proliferation through T6SS-dependent activation of the bone morphogenetic protein pathway. Cell Rep. 43, 113750 (2024).

79. Scopelliti, A. et al. Local control of intestinal stem cell homeostasis by enteroendocrine cells in the adult Drosophila midgut. Curr. Biol. CB 24, 1199–1211 (2014).

80. Guo, X., Lv, J. & Xi, R. The specification and function of enteroendocrine cells in Drosophila and mammals: a comparative review. FEBS J. 289, 4773–4796 (2022).

81. Beehler-Evans, R. & Micchelli, C. A. Generation of enteroendocrine cell diversity in midgut stem cell lineages. Dev. Camb. Engl. 142, 654–664 (2015).

82. Hudry, B., Khadayate, S. C, Miguel-Aliaga, I. The sexual identity of adult intestinal stem cells controls organ size and plasticity. Nature 530, 344–348 (2016).

83. Blackie, L. et al. The sex of organ geometry. Nature 630, 392–400 (2024).

84. Regan, J. C. et al. Sexual identity of enterocytes regulates autophagy to determine intestinal health, lifespan and responses to rapamycin. *Nat*. Aging 2, 1145–1158 (2022).

85. Regan, J. C. et al. Sex difference in pathology of the ageing gut mediates the greater response of female lifespan to dietary restriction. eLife 5, e10956 (2016).

